# Self-operated stimuli improve subsequent visual motion processing

**DOI:** 10.1101/2020.08.25.265918

**Authors:** Giulia Sedda, David J. Ostry, Vittorio Sanguineti, Silvio P. Sabatini

## Abstract

Proper interpretation of visual information requires capturing the structural regularities in the visual signal and this frequently occurs in conjunction with movement. Perceptual interpretation is complicated both by transient perceptual changes that accompany motor activity, and as found in audition and somatosensation, by more persistent changes that accompany the learning of new movements. Here we asked whether motor learning also results in sustained changes to visual perception. We designed a reaching task in which participants directly controlled the visual information they received, which we term self-operated stimuli. Specifically, they trained to make movements in a number of directions. Directional information was provided by the motion of an intrinsically ambiguous moving stimulus which was directly tied to motion of the hand. We find that movement training improves perception of coherent stimulus motion, and that changes in movement are correlated with the perceptual change. No perceptual changes are observed in passive observers even when they are provided with an explicit strategy to solve perceptual grouping. Comparison of empirical perceptual data with simulations based on a Bayesian generative model of motion perception suggests that movement training promotes the fine-tuning of the internal representation of stimulus geometry. These results emphasize the role of sensorimotor interaction in determining the persistent properties in space and time that define a percept.

## Introduction

Active interaction with the environment is a defining feature of our daily activities and critically relies on the interplay between motor and perceptual processes. This continuous exchange, besides promoting a proper calibration of sensory and motor systems and stabilizing the functional architecture in the respective circuits (1), allows mutual adaptation following sensory perturbations or motor training. The influence of movement on perception has been documented in situations in which movement changes – induced by adaptation or learning – elicit perceptual changes. For instance, adaptation to force fields (2–4), visuomotor rotations (5, 6) and optic prisms (7, 8) induce a shift in position sense. Somatosensory changes have also been reported as a consequence of motor skill learning, in which no perturbations were involved. For instance, training with maze tracing (9) or goal-directed movements (10) led to an increase in proprioceptive sensitivity. Presumably, as a result of the repeated pairing of somatic input and movement, training-related changes to movement and motor cortex are accompanied by somatosensory plasticity.

There has been substantially less work on the effects of motor learning on visual function. Most studies report transient changes to visual perception which accompany movement (see (11) for review). Movement execution (12), movement planning (13) and cognitive expectations (14) each shape the visual perception of a moving stimulus. Movement can bias perceptual sensitivity toward visual events that either share features with what we are currently doing (15) or that deviate from the expected sensory consequences of our movements (12). This suggests that action may guide inferential processes from visual cues to categories and suggests that cognitive or context expectations can concurrently influence our perceptual judgements. Yet, these studies suggest no evidence of sustained effects that reflect visual perceptual learning, as an enhancement of perceptual discrimination/detection capabilities *following* motor practice with a visual stimulus. Vision is indeed a highly reliable source of information and it is difficult to induce changes in visual perception, at least when simple forms of visual feedback like displayed positions or trajectories are involved. However, vision, like other exteroceptive senses, does not provide a unique interpretation of reality, for example when we look at objects with shadows or in different lighting. During development, through active interaction with the environment we learn to combine different cues and contextual information to find a unique solution that usually corresponds to veridical interpretation. A grating that moves through an aperture is an example of an inherently ambiguous visual pattern, as its movement direction cannot be uniquely determined from visual information alone (16–18). Moreover, it has the desired feature of selectively activating specific early spatiotemporal frequency channels in the cortex. When we observe two superimposed gratings moving in different directions – a ‘plaid’ stimulus – we tend to integrate their drifting speeds into one coherent motion. Alternatively, the plaid can be perceived as two separate gratings, which slide over each other in different directions – a situation referred to as transparent motion (19). By varying the features of the individual gratings the perceptual ambiguity can be manipulated (20–22). As a general rule, when the two component gratings are more balanced, i.e. they are more similar in terms of spatial frequency, contrast, and luminance, the plaid is more likely perceived as a coherent pattern moving in one direction.

To understand how movement affects the way we make sense of this complex visual information, we specifically ask if experiencing the visual consequences of self-generated movements can promote persistent perceptual changes that affect subsequent judgement tasks. In this respect, the point is not learning a motor skill or adapting to external perturbations, but just exercising sensorimotor contingencies, by experiencing the sensory consequences of self-generated movements. Accordingly, we designed a motor task in which the direction and speed of the hand is continuously displayed as a plaid moving through an aperture. We then looked at whether motor training affects the ability to perceive subsequent plaid motions. A Bayesian generative model of the perceptual process helped to identify the underlying mechanisms.

## Results

The experimental apparatus and procedure used in this study are illustrated in Figure 1. Figure 1b shows the plaid stimulus, which is formed by two gratings with different orientations. When a single moving grating is observed through an aperture, only the component velocity perpendicular to its orientation can be perceived. By adjusting the relative difference in the contrast of the two gratings, they appear either to be sliding one over the other in directions ***υ***_1_ and ***υ***_2_ or as a single plaid pattern moving in direction ***υ***. In this way, the extent to which one perceives the coherent motion of a single plaid or two separate gratings can be manipulated.

**Fig. 1.**
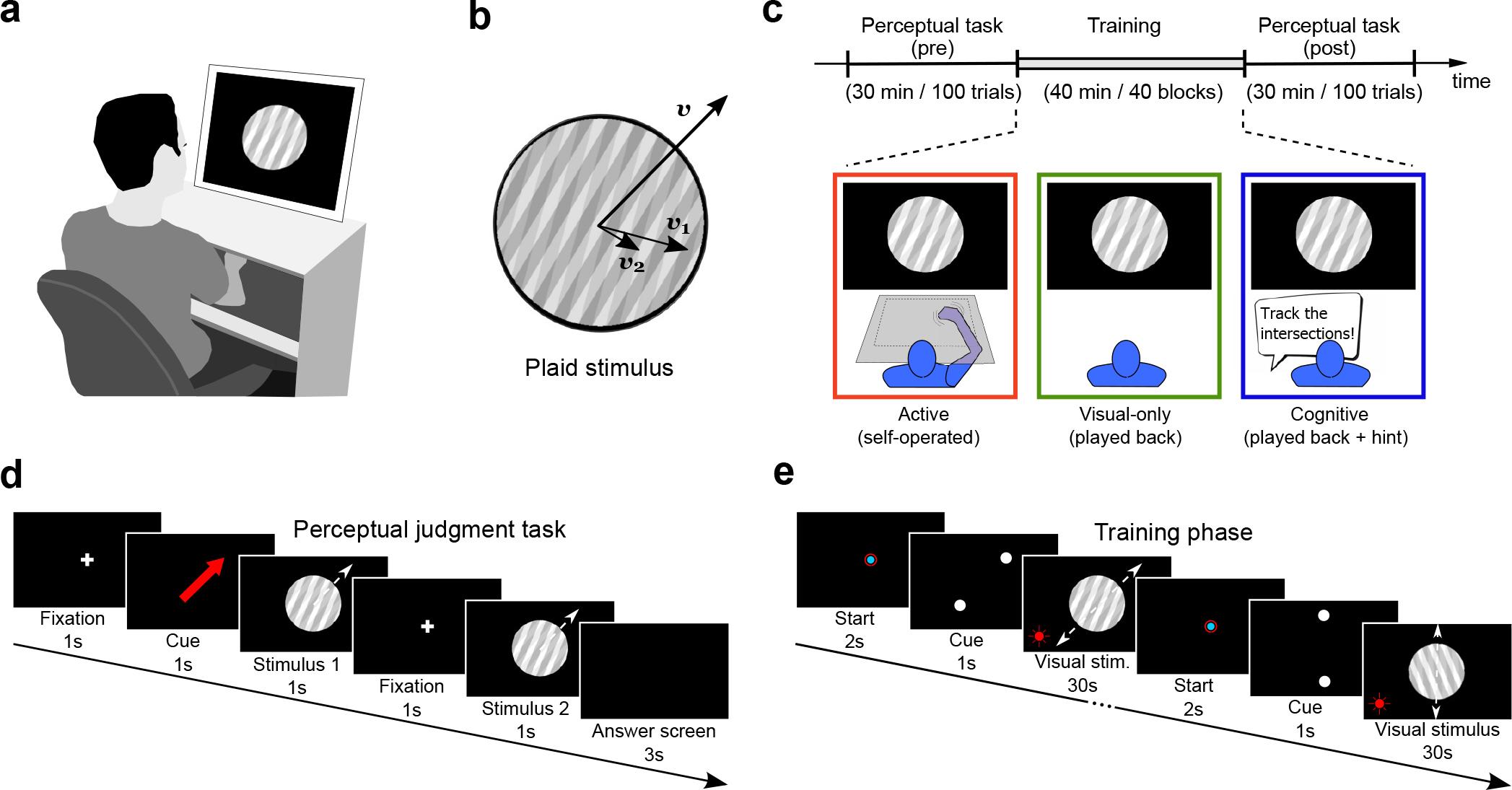
Experimental setup and protocol. a) Experimental setup: The participant is seated in front of a screen and is exposed to moving visual stimuli (plaid). During active training they perform planar movements which result in motion of the plaid on the screen. Visual feedback of the arm is blocked. b) A plaid stimulus with velocity *υ*, composed of two gratings moving at velocities *υ*_1_, *υ*_2_. c) The experimental protocol has three phases: Participants start with a perceptual judgement task, then they perform a training task, and finally they repeat the perceptual task. Participants were divided in three groups, each with a different training condition: active, visual-only, and cognitive. d) The perceptual task is a 2AFC paradigm. Participants see two consecutive moving plaid stimuli, and are asked to choose which stimulus is moving in a direction more similar to that of the red arrow. e) During training participants are exposed to moving plaids. In the active group they perform planar hand movements to control the plaid motion on the screen, while participants of both the visual-only and cognitive groups observe played-back motions. In the cognitive condition, participants are instructed to track the intersections of the gratings.

In the experiment, participants undergo an initial perceptual judgement task to assess perception of plaid motion for different contrast values (see Figure 1d). The perceptual task involves a two-alternative forced-choice (2AFC). Participants are presented with two consecutive moving plaids, which differ in the amount of the contrast difference. They are required to indicate which plaid is moving in a direction most similar to that shown by a red arrow. One of the two plaids, a Reference stimulus, has a fixed contrast difference Δ*c_R_* between the two gratings. In the other, a Test stimulus, the contrast difference Δ*c_T_* is systematically varied, and it is always less, which makes it easier to detect the plaid motion direction. This is followed by a training phase (see Figure 1e), after which the perceptual task is repeated. In all conditions the plaid motion is seen through an aperture. Three different groups of participants were tested. In an active training condition participants use self-operated plaids: they control the plaid motion by moving their hand, such that the direction and velocity of the moving plaid corresponds to that of the hand; vision of the hand is blocked. Participants were instructed to make continuous movements back and forth between two circles that were presented briefly at the start of a continuous movement trial. The contrast difference Δ*c* between the single gratings that form the plaid is based on the individual threshold estimated from the pre-training perceptual task. In a visual-only condition the participant sees a played-back moving plaid stimulus of another participant. In a cognitive condition the stimulus is identical to that in the visual-only condition, and in addition the experimenter instructs the participant to track the intersections of the two gratings. The motion of the intersections corresponds to that of the plaid. This provides participants with an explicit strategy that enables them to correctly estimate the plaid motion.

### Perceptual learning

The results of perceptual task (the probability of selecting the Test stimulus as a function of the contrast difference Δ*c_T_*) and, in particular, training related changes in perception are presented in Figure 2a. Both threshold differences (ΔTh = Th_post_ – Th_pre_) and differences (ΔSlope = Slope_post_ – Slope_pre_) in the slope of the psychometric function are shown. Better perceptual performance is reflected in an ability to select the Test stimulus under conditions of greater contrast difference, that is for larger values of Δ*c_T_*. Both threshold and slope values were estimated using the adaptive Ψ procedure (see Methods). It is worth noting that the number of trials (100) chosen for the perceptual judgment task allows full convergence for perceptual threshold values, but not for the slope estimates (see Methods) (23). The values of the perceptual slope are shown for completeness and to allow for qualitative analysis of the results. Figure 2b shows psychometric threshold differences and slope differences for all participants in each experimental condition. It can be seen that there are changes in the psychometric threshold for the active group, only, and no changes in slope in any of the experimental conditions. Statistical analyses were conducted using difference scores which were found to be normally distributed (P > 0.05; Anderson-Darling test), whereas the pre-training and post-training perceptual values were not normally distributed (P <0.05). We ran nonparametric tests (Kruskal-Wallis) to verify that baseline values for threshold and slope did not differ. We tested for differences in both the threshold (ΔTh) and the slope (ΔSlope) of the psychometric curves. We observed a highly significant difference in threshold between experimental conditions (P =0.0002; F(2,27)=7.45; One-way ANOVA), and no reliable difference in slope. Post-hoc analyses (Bonferroni-Holm) revealed a significant difference in the threshold of the active and visual-only conditions (P = 0.003) and between the active and cognitive conditions (P = 0.019).

**Fig. 2.**
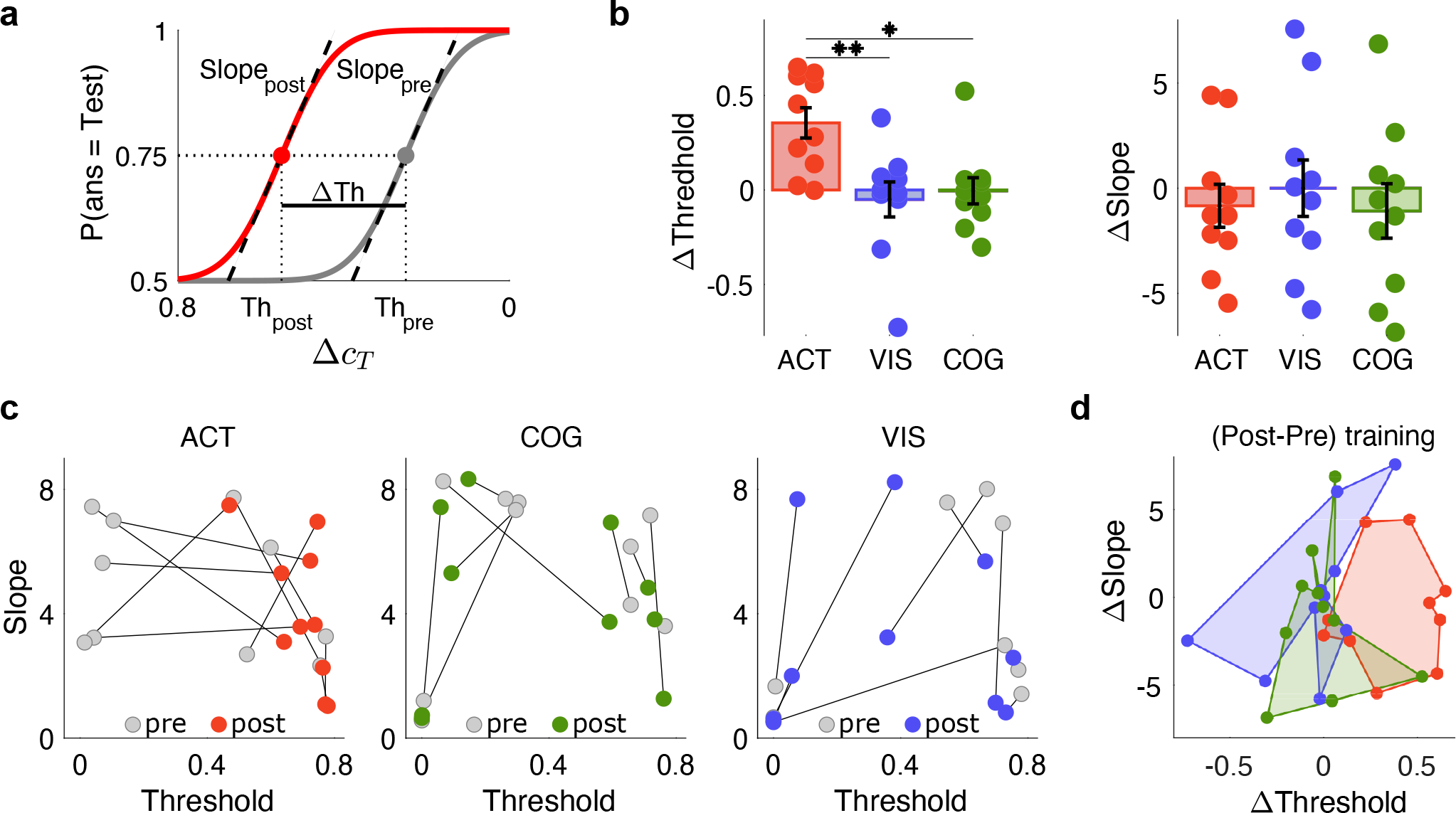
Results of the perceptual judgement task. a) Representative psychometric curves. Each curve shows the probability that the participant chooses the Test stimulus over a range of relative contrast differences Δ*c_T_*. Grey curve represents the perceptual baseline of a representative subject (pre-training), whereas the colored curve indicates the perceptual change (post-training). Solid lines represent the average values, the filled circles indicate the 75% threshold value (Th_pre_, Th_post_), and the dashed black lines show the slope of the curves at the threshold point (Slope_pre_, Slope_post_). The horizontal black segment displays the threshold difference, ΔTh = Th_post_ – Th_pre_. b) Colored bars represent the average values of threshold differences ΔTh and slope differences ΔSlope in all three experimental conditions. Dots represent the individual values for each subject. Error bars denote standard errors. The average value of ΔTh in the active group is significantly greater than in the visual-only (P = 0.003) and the cognitive (P = 0.019) groups. c) Qualitative analysis of inter-subject variability is shown in terms of threshold and slope changes for the individual subjects in each condition. In all three conditions, the grey dots represent the pre-training values. d) Qualitative analysis of inter-subjects variability is shown in terms of the minimum polygons enclosing all data points in each group (active: red; visual-only: blue; cognitive: green).

Figure 2c,d summarizes inter-subject variability. After training, all subjects in the active group exhibit a threshold value which is close to the maximum value of 0.8, in other words they correctly select the Test stimulus throughout the entire range of Δ*c_T_*. In contrast, subjects in the visual and cognitive groups exhibit no consistent trend in either threshold or slope. Figure 2d displays inter-subject variability in terms of the minimum polygons enclosing all data points in each group. Before training, the subjects within each group exhibit a similar amount of variability. After training, the subjects in the active group display a polarization towards greater threshold values. The distribution of the differences between ‘pre’ and ‘post’ values in both thresholds and slopes shows for the three training conditions a different clustering in three distinct regions of the ΔThreshold —ΔSlope plane.

### Motor training

During movements, the subjects in the active group initially exhibit a positive (counterclockwise) directional bias; see Figure 3a. This motor bias tends to decrease with training. We observed a highly reliable correlation (R^2^ =0.79, P = 0.002) between the change in movement direction (ΔMotor bias) observed during training and the perceptual change before and after training (ΔPerceptual Threshold); see Figure 3c. Figure 3b summarizes the inter-subject variability in both perceptual and motor performance. Individuals exhibiting a lower initial perceptual threshold (hence a ‘poor’ sensory performance) benefit more from motor training. Subjects with an initially higher perceptual thresholds (i.e. an already good sensory performance) exhibit a lower motor bias; in this case the perceptual thresholds remain constant or increase.

**Fig. 3.**
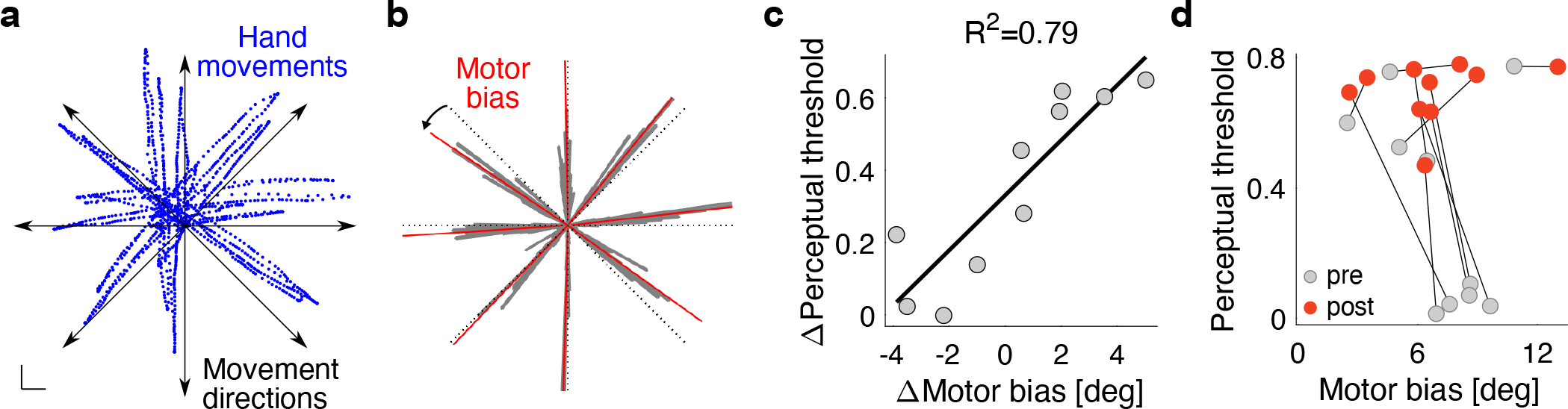
Results of active motor training. a) For a typical subject, the trajectories (blue dots) of hand movements across all trials of the first block for all movement directions are represented. Solid black lines represent the tested directions. Scale bar: 2cm. b) For the same typical subject, the frequency distribution (grey polar histogram) of hand movement across all trials of the first block for all movement directions are represented. Dashed black lines represent the tested directions. Red solid lines represent the median value of the motor bias for each direction. The black arrow shows the sign of the directional bias. Bin size: 1 deg. c) Relationship between perceptual change from before to after training (ΔPerceptual threshold), and the change in movement direction (ΔMotor bias) that occurred in conjunction with training (P = 0.0006). d) Qualitative analysis of inter-subject variability is shown in terms of perceptual threshold and motor bias changes for the individual subjects in the active group.

### Computational model

A Bayesian framework was used to model the way humans perceive plaid motion. The model incorporates a number of empirical findings on how the perception of single gratings is affected by contrast.

We specifically assumed that perception is affected by both random and systematic effects. In particular, the variance in the perceived velocity of a grating (here after referred to as perceptual variance) increases with the negative power of the contrast (24), where *s*^2^ is the perceptual variance at maximum contrast, and *q* is the power exponent. We also assumed a reciprocal influence of one grating on the perception of the other grating’s velocity (cross-talk), proportional to their contrast imbalance through a parameter *k*. Finally, we assumed a Gaussian prior for grating velocities with zero mean and variance *σ_p_*^2^. As a whole, the model is fully characterized by the parameter vector *w* = [*s*^2^, *q, k, σ_p_*^2^]^*T*^; see Methods for further details.

Unlike previous Bayesian formulations (25, 26), this model is able to reproduce key findings concerning the directional bias in the perception of plaid movements (27, 28); see the SI Appendix for details.

The perceptual judgement task was modelled as a binary decision between two possible alternatives, Test (T) or Reference (R). The probability of choosing T as a function of the estimate of plaid direction, 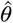, and the contrast difference Δ*c_T_* in the test stimuli, i.e. 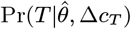, is modelled as a Bayesian decision process (see Methods, Eq. 6). The predicted psychometric curve is a function of the model parameters *w*, i.e. 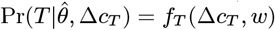, see Figure 4.

**Fig. 4.**
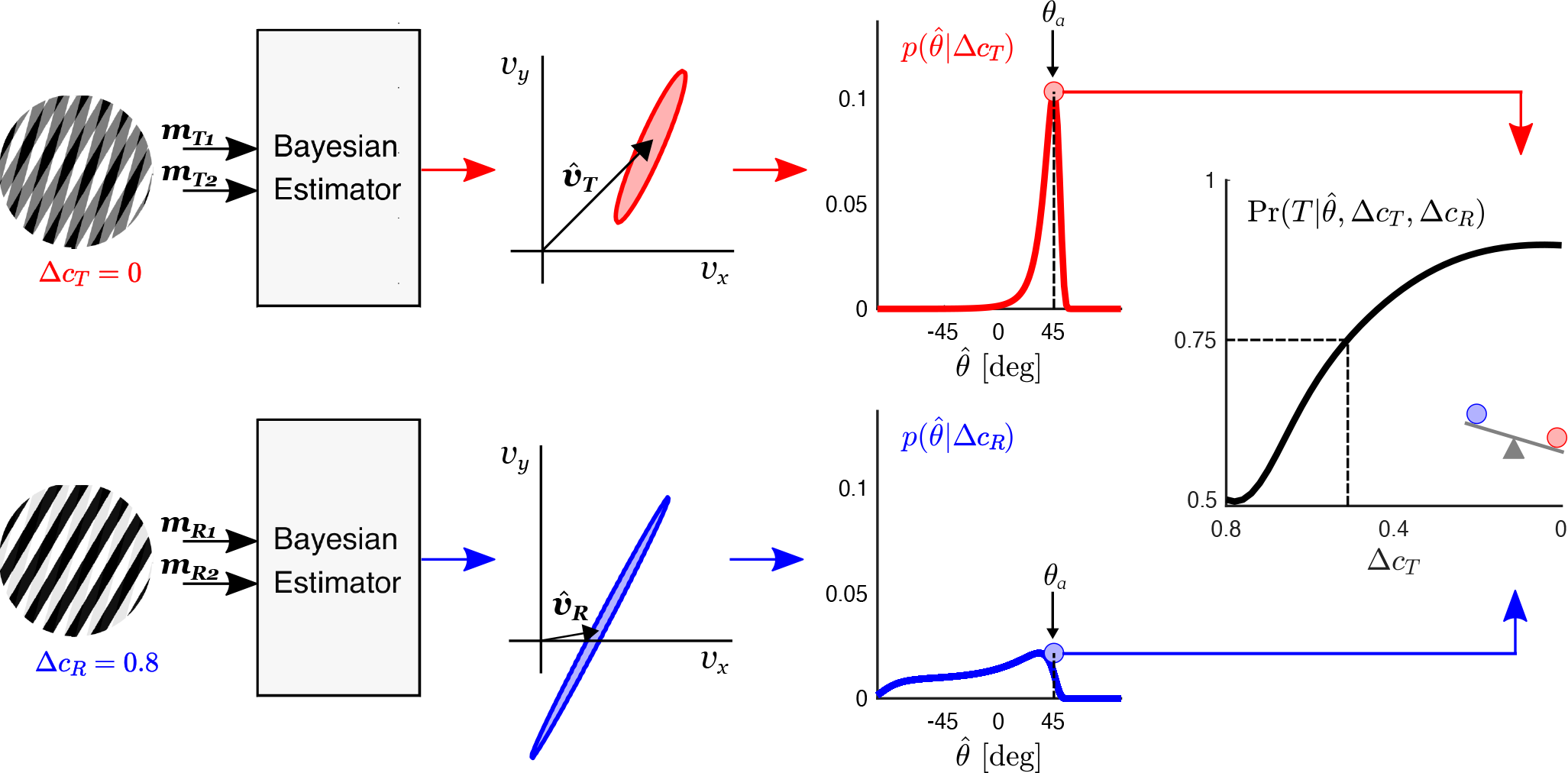
Bayesian model for the plaid estimation process and forced-choice paradigm. A Test (T) and Reference (R) plaids are shown, with Δ*c_T_* = 0 (top) and Δ*c_R_* = 0.08 (bottom). For each plaid, the optimal estimate of plaid velocity, 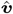, is represented. 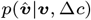 has a normal distribution, in which both mean and covariance depend on the relative contrast difference, Δ*c*. The probability of estimating a plaid direction 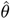 given a specific Δ*c* is given by 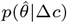. The psychometric curve represents the probability of answering T as a function of the contrast difference Δ*c_T_* and Δ*c_R_*, i.e. 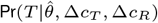, where 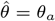 (45 deg in our experiment).

We fitted the model to the data and identified the model parameters *w* before and after each of the training conditions. Figure 5 summarizes the fitting results. Figure 5a shows the average value of each model parameter among subjects, before (grey boxes) and after (colored boxes) each training condition. We found a significant change in 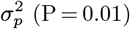, *k* (P = 0.026), and *s*^2^ (P = 0.025) parameters in the active group alone. Perceptual threshold values for the model are estimated at the 75th percentile (see Methods) and show a high correlation with the threshold values in the experimental tests (see Figure 5b). Figure 5c shows psychometric threshold differences (ΔTh = Th_post_ – Th_pre_) calculated from model fitting for all the participants in each experimental condition (see Eq. 6). It can be seen that as in the empirical results there are significant changes in the psychometric thresholds in the active group alone. These observations are confirmed by statistical analysis. We tested for differences in threshold (ΔTh) of the psychometric curves. We observed a significant difference in threshold between experimental conditions (F(2,27) = 7.76; P = 0.002). Post-hoc analyses (Bonferroni-Holm) revealed a significant difference in the threshold of the active and visual-only conditions (P = 0.003) and between the active and cognitive conditions (P = 0.024). These results on the perceptual threshold of the curves obtained by fitting the data with the Bayesian generative model are in agreement with those found in the results of the perceptual judgement task by fitting the data with the cumulative Gaussian function, as shown in Figure 5c; see Methods. Moreover, we found a highly reliable relationship between the change in the model parameter *k* (Δ*k*) from before to after training, and the perceptual change from before to after training (ΔTh) (R^2^ = 0.81, P = 0.0004); see Figure 5d. This means that for subjects with greater perceptual changes the model predicts a greater decrease in the cross-talk parameter *k*.

**Fig. 5.**
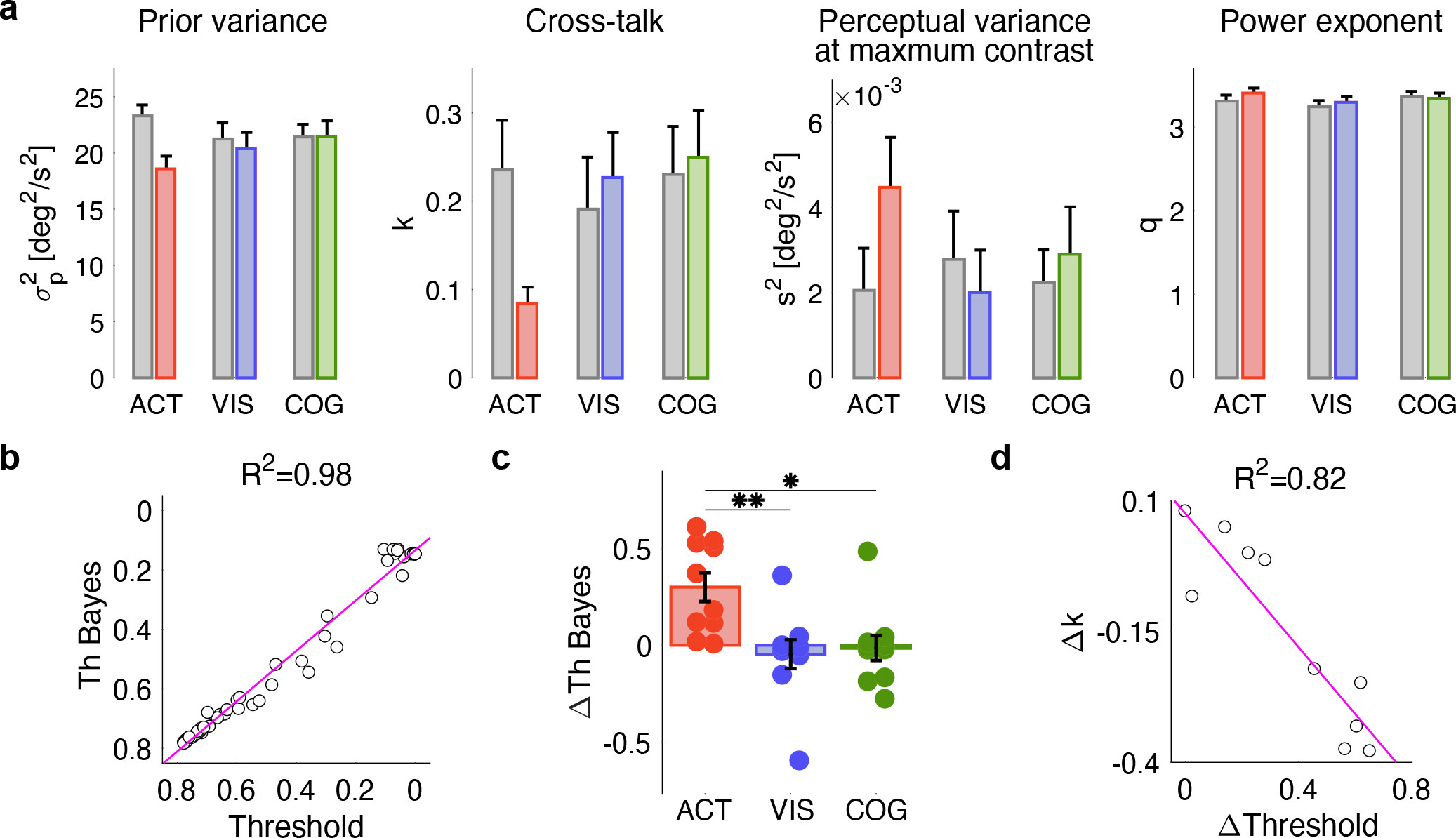
Results of model fitting. a) Comparative parameter fitting for the different training conditions: 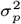 (Anderson-Darling test for normality: P > 0.05, T-test P = 0.01 for the active group), *k* (Anderson-Darling test for normality: P > 0.05, T-test P = 0.026 for the active group), *s*^2^ (Anderson-Darling test for normality: P = 0.003, Wilcoxon rank sum test P = 0.025 for the active group), and *q*. Grey and colored boxes refer to pre- and post-training conditions, respectively. b) Correlation between thresholds estimated using perceptual test data and from the Bayesian generative model (P = 0.001). c) Average thresholds of Bayes’ model psychometric curves over all subjects of each group. d) Correlation between the perceptual threshold change, ΔThreshold, and the corresponding variation of the cross-talk parameter, Δ*k*, (P = 0.004).

## Discussion

The present study shows that active interaction with an ambiguous visual stimulus alters the subsequent perception of stimulus motion. Three groups of participants performed the same perceptual task before and after training. Self-operated motion of plaid stimulus was generated by an active group that performed planar movements. This was designed to assess whether perceptual decisions regarding plaid movements were affected by actively interacting with the stimulus. A visual-only group observed played-back plaid motion that was generated by another subject. This condition quantified the effect of prolonged exposure to the moving plaid stimulus. A cognitive group experienced the same stimuli as the visual-only group. These subjects were additionally instructed to track the gratings intersections and thus had an explicit strategy which would enable them to follow the coherent plaid motion. We found that the perceptual threshold for the direction of plaid motion changed significantly following training only in the active movement condition, where it showed more robust perceptual integration against contrast imbalance in the plaid. There were also practice related changes to movement. Movement direction changed over the course of training presumably because the plaid, which effectively serves as a cursor showing movement direction, is more easily seen by subjects as moving in the remembered target direction. We found that the change in perceptual threshold was strongly correlated with the change in movement direction measured during the active training, consistent with the idea that the perceptual change is tied to motor training. A quantitative model suggests that movement training affected perceptual judgement by improving the accuracy of the internal representation of the plaid geometry. The findings indicate that motor training resolves visual perceptual ambiguity and contributes to changes in visual perceptual ability.

### Motor training implications for perceptual discrimination

A small number of studies have examined the effects of motor learning on vision. Brown et al. (29) found that movement initiation toward a moving object, that was to be intercepted, differed depending on the direction of a previously learned force field, indicating that expectations regarding visual motion are altered as a result of learning. Beets et al. (30) showed that when participants trained to make movements that violated to different degrees the 2/3 power law, there were improvements in visual discrimination of movements that corresponded to those which they experienced during training. These studies indicate that motor learning can induce a bias in visual perception. The present study suggests that movement training plays an even more pivotal role. Indeed, visual perception is inherently ambiguous. Movement training leads to a reduction in perceptual uncertainty, and to a change of perceptual sensitivity which in the present case is related to a stimulus parameter (gratings’ contrast difference) that is not directly controlled during training. Both movement training and perceptual change occur here without feedback motor error provided to participants during training. Changes in movement direction within single training blocks and over the entire training session are significantly correlated with the observed perceptual change (between pre- and post-training perceptual tasks) suggesting that the two kinds of learning are cross-related. If we assume that visual motion perception is based upon an empirical strategy which serves to resolve perceptual uncertainty (31, 32), the perceived plaid motion direction is determined by accumulated sensorimotor experience. Perceptual decisions regarding motion direction provide, in turn, sensory evidence that instructs behaviour. Our results suggest that visual function over time can be adapted with training, which is provided by interaction with the stimulus.

### Modeling the fine-tuning of internal representation of plaid geometry

Several studies (25, 26) used a Bayesian framework to model the perceptual task of estimating the velocity of the plaid from the perceived velocities of the two gratings. These models posit that prior information and an internal (neural) representation of plaid geometry are combined to obtain the expected value of plaid velocity (26). Prior information captures the participant’s prior experience with observing moving patterns and is summarized by the statistical distribution of the plaid velocities. The representation of plaid geometry approximates the mapping from plaid to gratings’ velocities. Accordingly, inaccurate perception of plaid motion may be due to (i) an inaccurate representation of plaid geometry (sensory model), (ii) an inaccurate perception of the velocity of each grating (noise variance), (iii) the bias introduced by previously experienced plaid motions (the prior); or a combination of the above.

Recent studies suggest that this Bayesian formulation cannot account for key observations in the way the perception of plaid direction is affected by speed (28). In the present paper, we made a number of specific assumptions on how the gratings’ contrasts affect the perceived plaid velocity. First, consistent with previous findings (24) we assumed that the variance of sensory noise in perceiving the velocity of a grating is proportional to an inverse power of the grating contrast. This is reflected in two model parameters: the variance at maximum contrast and the power exponent. With these simple additions our model predicts those same observations (28) that have been claimed to falsify Bayesian models of plaid perception. We also posited an additional effect in the representation of plaid geometry: if two gratings have different contrasts, the represented direction of a grating affects that of the other – this was denoted by the ‘cross-talk’ parameter (cf. (27)). This effect results in a systematic error in the representation of plaid geometry.

For each participant we estimated the parameters that maximise the model likelihood given the data from the perceptual judgement task. For each experimental condition, we then assessed the model parameter changes from before to after training. Significant changes in the model parameters were obtained for the active training condition alone. Specifically, we found that participants in this condition exhibited a significant decrease in the cross-talk parameter and an increase of the power law exponent. The reduction in cross-talk leads to a more accurate representation of the direction of the gratings and therefore a more accurate representation of plaid geometry. The increase of the power law exponent leads to a decreased sensitivity of sensory noise to contrast.

Note that the cross-talk decrease exhibits a strong correlation with the observed change in perceptual threshold. That is, participants who exhibit greater perceptual changes also show a greater reduction in the cross-talk. This finding suggests that motor training in this task leads to a fine-tuning of the internal representation of plaid geometry.

Why does movement improve the sensory model while observation on its own does not? One possible explanation is that during movement, the sensory model predicts the sensory consequences of movements – the expected movements of the gratings. The mismatch between these predictions and the observed movements of the gratings – sometimes called sensory prediction error – is the source of information which can be used to adapt the sensory model. This information is not available during passive observation of plaid movements. Consistent with this view, sensorimotor adaptation to dynamic or visual perturbations has been reported to critically depend on the availability of a sensory prediction error signal (33, 34).

### Implications for neural representations of complex visual motion

As plaid stimuli are composed of a minimal number of one-dimensional Fourier components (two), each selectively recruiting narrow early vision oriented band-pass frequency channels, they can contribute to understanding how these channels are involved in the perceptual learning of coherent sensorimotor dependencies.

Finding a solution to the plaid motion problem can be related to the evidence from component-motion and pattern-motion cells, observed respectively in striate and extrastriate areas along the primary visual motion pathway, such as area V3A and middle temporal area (MT or V5) (35). Ultimately, the steps in the formation of perceptual decisions and/or guidance of visual behaviours can be linked to higher-level brain areas (e.g., lateral intraparietal cortex and prefrontal cortex), which are often described as “evidence accumulators” (36–38). The present model simulation suggests a reduction, after training, of the cross-talk between the two gratings in the corresponding sensory channels, when the gratings contrasts are unbalanced, as well as a reduction of the noise variance. The decrease in cross-talk magnitude would be consistent with an early neural instantiation of the perceptual learning process, which might occur at the coding stage of the component motion directions (cf., contrast normalization processes in V1). On the other hand, since the cross-talk in the model acts on gratings’ velocities, it requires a pooling of the responses of different oriented channels, and its change might occur at pattern motion coding in an extrastriate area. Notably, the null effect of visual-only training leads us to exclude a role of a oculomotor-specific, but rather a reach-specific sensorimotor cortical area. Specific experiments and recordings of neural correlates would be necessary to disambiguate the different hypotheses. From a broader perspective, this study suggests a shift in focus to pattern and motion vision investigation, which includes continuous interaction with visual stimulation.

## Materials and Methods

### Subjects

A total of 30 subjects (11 male and 19 female, 18–30 years old) participated in this study. All participants had normal or corrected-to-normal vision and reported no history of neurological disorder. They were naïve to the purpose of the study and received written and verbal instructions before the start of the experiment. Each participant was randomly assigned to one of three groups (10 participants per each group). The research conforms to the ethical standards laid down in the 1964 Declaration of Helsinki that protects research participants and was approved by the Ethical Committee of the Dept of Informatics, Bioengineering, Robotics and Systems Engineering, University of Genoa. Each subject signed a consent form conforming to these guidelines.

### Apparatus

Visual stimuli were presented on a 19-inch LCD monitor (Samsung B2430L) at 1920× 1080pixels, and refreshed at 60 Hz. In a dimly lit room, participants were seated in front of the screen at about 57 cm of distance, so that the visual angle of the whole display was 60deg; see Figure 1a. In one part of the experiment (see below) participants grasped the puck of a digitizing tablet (CalComp, Inc, 3200-series DrawingSlate II, Model 32120) to actively drive the motion of the visual stimulus using planar movements. The digitizer had a 305 mm×457 mm workspace, and a 125 Hz sampling rate. The center point of the screen was mapped onto the center of the digitizing tablet, with a 1:1 scale factor, see Figure 1c.

### Stimuli

We presented a plaid stimulus composed of two square-wave gratings through a circular aperture, about 13deg in diameter, on a black background, as shown in Figure 1a. The luminance of the black background outside the aperture was 0cd/m^2^. The two gratings had normal directions *θ*_1_ and *θ*_2_. The plaid moved at speed *v* = 5deg/s in the direction *θ* = 45deg (from the lower left corner of the screen to the upper right corner). Δ*θ*_1_ = *θ*_1_ – *θ* = −60deg and Δ*θ*_2_ = *θ*_2_ – *θ* = −75.5deg define the relative directions of the individual gratings with respect to the direction of the plaid. With this geometric arrangement the ratio between the two gratings speeds is cosΔ*θ*_1_/ cosΔ*θ*_2_. In particular, stimuli were designed as plaids, whose direction fell outside the range of the directions of the two component gratings (type II plaid, see (39)). In particular, we chose gratings directions that were relatively close to one another, and far away from the direction of the whole plaid pattern. Because of this geometric arrangement the plaid motion direction was distinct from that of the gratings (40), and the directions of gratings were sufficiently close to one another to be interchangeable with their average. Each grating was composed of dark (55-65cd/m^2^) and light (115-125cd/m^2^) stripes, and a spatial frequency of 0.6cycle/deg. Stimulus was presented in transparency (19, 22, 27), and the perceptual uncertainty was modulated by varying the contrast level of each grating. The overall plaid image was defined as:

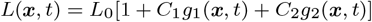

where *L*_0_ is the mean intensity, *g*_1_ and *g*_2_ are the functions that defined the two component gratings, and *C*_1_ and *C*_2_ are the gratings’ contrast levels, respectively (19). The total contrast *C* = *C*_i_ + *C*_2_ was kept constant, and the relative contrast difference between the gratings of each plaid was defined as Δ*c* = |*C*_1_ – *C*_2_|/*C*. In all experiments we set *L*_0_ ≃ 90cd/m^2^ and *C* = 0.5. Participants were instructed to maintain fixation at the center of the stimulus during the entire duration of stimulus presentation. Stimuli were generated using Psychophysics Toolbox for Matlab (41, 42).

### Experimental Protocol

The experimental procedure had three phases; see Figure 1c (top). Participants were initially administered a perceptual judgement task (pre-training test). Next, they underwent a training phase under a variety of conditions (see below). After training, they repeated the perceptual judgement task (post-training test).

### Perceptual judgement task

The purpose of this test was to quantify the ability to correctly assess the direction of plaid motion as the relative contrast difference of the two gratings (Δ*c*) was varied. The test used a 2-alternative forced-choice (2AFC) paradigm; see Figure 1d. Each trial started with a fixation point (black screen with a white cross at the center) displayed for 2 s. Then a red arrow with a *θ_a_* = 45deg direction, was displayed for 1 s. Finally, two different plaids were presented for 1 s each, separated by a 1 s fixation point. The two plaids were identical and both moved in the direction *θ* = 45 deg, but had different Δ*c*. At the end of the trial, participants were asked to choose which of the two plaids had a movement direction which was most similar to that denoted by the red arrow. They had to provide an answer by pressing the left or right arrow on the keyboard within a 3s time limit to indicate the first or the second plaid, respectively. Throughout the entire test, one plaid (Reference stimulus, R) had a constant contrast difference, Δ*c_R_* = 0.8, which corresponds to a large imbalance in the contrast of the component gratings, which in turn favours the perception of the individual gratings motions. In the other plaid (Test stimulus, T), the contrast difference Δ*c_T_* changed on each trial, within a 0-0.8 range. The Test and Reference plaids were presented in random order.

We used a Bayesian adaptive procedure – Ψ (Psi) method (23, 43) – to select the value of Δ*c_T_* on the current trial, based on the participant’s answers in the previous trials. We took the selection of the Test stimulus as the correct answer. Every time the subject answered correctly, the Δ*c_T_* value was increased, so that it gradually became more and more similar to Δ*c_R_*.

The entire perceptual judgment test took a total of 100 trials to complete, which corresponded to a duration of about 30 min.

### Active motor training

Participants were instructed to perform out and back planar arm movements between two briefly presented visual cues, in a target direction *θ_T_*; see Figure 1c (bottom, left). The motion of a plaid on the screen was continuously yoked to the instantaneous direction of hand movement, *θ*(*t*), so that the two gratings moved in directions *θ*_1_(*t*) = *θ*(*t*) + Δ*θ*_1_ and *θ*_2_(*t*) = *θ*(*t*) + Δ*θ*_2_ while their relative orientations with respect to plaid motion, i.e. Δ*θ*_1_ and Δ*θ*_2_ remained constant.

The training phase was organised into a series of trials, each characterized by a different target hand direction.

At the beginning of each trial, participants had to place the hand (depicted as a blue cursor on the screen) inside a start region (circle on a black background) and hold it there for 2 s. Then both the start region and the cursor disappeared, and a circular aperture was displayed. Two white circles placed just outside the aperture, were displayed for 1 s, at opposite sides with respect to the center of the aperture, 28deg of visual angle from one another with respect to the participant. The circles indicated the target hand direction for that trial. As the circles disappeared, a plaid appeared inside the aperture. Participants were instructed to move the hand back and forth in the target direction, between the two remembered circle positions. Participants were encouraged to maintain a speed no greater than 5deg/s – the speed of the plaid used in the perceptual judgement task. To aid in maintaining the correct speed, participants continuously received visual feedback on movement speed (circular spot in the bottom left corner of the screen; green if the speed was ≤ 5deg/s, red otherwise). Each trial had a fixed duration of 30 s.

During training, participants were prevented from seeing their arm, so that the only visual information about their movement direction was provided by the plaid motion. During the movement training phase, the relative contrast difference Δ*c* in the plaid was set to that subject’s threshold level, as estimated at the end of the pre-training perceptual judgement task. The entire training protocol involved four target directions (0 deg, 45 deg, 90 deg, 135 deg) each repeated 10 times in pseudo-random order, for a total of 40 trials and an approximate duration of 40 min.

### Visual-only training

Participants were instructed to observe on the screen a plaid moving through an aperture, while performing no movements. The plaid stimulus was the playback of a stimulus generated by another participant in the active training group; see Figure 1c (bottom, middle). Again, the total duration of this phase was about 40 min.

### Cognitive training

As in the visual-only training condition, participants had to observe on the screen a plaid while performing no movements. In addition, they were provided a hint to estimate the plaid movement direction – track the movements of the grating intersection points which have the same direction and speed of the plaid; see Figure 1c (bottom, right). Again, the total duration was about 40 min.

### Data analysis

For each subject, we quantified performance in the perceptual judgement tasks before and after training by estimating a psychometric curve using a Bayesian adaptive Ψ (psi) method (23, 43) and assuming a normal cumulative distribution function. We used the threshold and slope of the estimated psychometric curve as measures of perceptual performance. The threshold value is defined as the Δ*_c_T__* value corresponding to 75% probability of selecting the Test stimulus, whereas the slope is defined as the inclination value of the psychometric curve at the threshold point. It is important to note that the number of trials (i.e. 100) chosen for the perceptual judgement task allows full convergence for the perceptual threshold values, but not for the slope estimates (23). We then assessed whether perceptual performance was affected by training in the active, visual or cognitive training conditions. To do this, we took perceptual threshold and slope before training (Th_pre_, Slope_pre_) as the baseline perceptual performance. We then looked at the threshold and slope after training (Th_post_, Slope_post_). For each quantity and for all experimental conditions, we first assessed normality (Anderson-Darling test). If normality was not ruled out for perceptual thresholds and/or slopes, we ran a repeated measures 2-way ANOVA with time (PRE, POST) and experimental condition (active, visual, cognitive) as within- and between-subject factors.

In case the normality assumption had to be rejected, we used a non-parametric test (Kruskal-Wallis) to assess differences among conditions in the perceptual baseline (Thpre and Slopepre). We then focused on the training-related change (ΔTh = Th_post_ – Th_pre_; same for slope). We tested for differences among experimental conditions, using 1-way ANOVA if normality was not ruled out; a non-parametric test (Wilcoxon Rank Sum) otherwise. Post-hoc analyses were conducted using pairwise t-test, with a Bonferroni-Holm correction.

Finally, we examined movements of the hand in the active motor training condition. For each trial, we calculated the statistical distribution of hand velocities (direction and magnitude), by separately accounting for forward and backward movements. We subtracted the target direction from the distribution of movement directions and then took the mean (bias) and standard deviation of the directional error for each block and each subject. We assessed how these quantities changed over the course of training (correlation with block number) and whether these changes correlated with changes in perceptual performance.

### Computational Model

#### Plaid geometry

Plaid geometry is completely specified by the overall plaid velocity, ***v*** and by the directions of the two gratings, *θ*_1_ and *θ*_2_. The velocity of one single grating, *v_i_, i* = 1, 2 is calculated as the projection of the plaid velocity onto the grating’s normal direction: 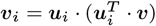 where ***u***_*i*_ = [cos *θ_i_* sin*θ_i_*]^*T*^, *i* = 1, 2; see Figure 1b. The above expression can be rewritten as

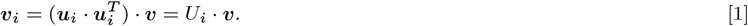

#### Sensory system

We assumed that the perceived velocity of each grating, ***m***_*i*_, *i* = 1, 2, is affected by additive zero-mean Gaussian noise, so that:

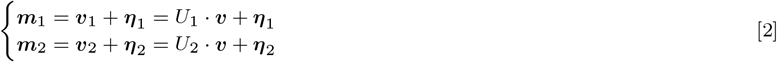

where ***η***_*i*_ ~ Normal(0, *Q_i_*), *i* = 1, 2 and the noise covariance matrix, *Q_i_*, is defined as

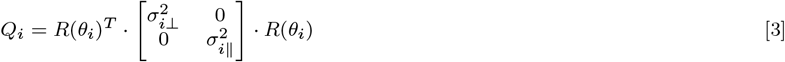

where 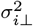 and 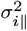 are the noise variances in directions that are perpendicular and parallel to grating *i*, and *R*(*θ_i_*) is a rotation matrix. As in (26), we set 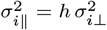 with *h* = 0.3, so that the covariance matrix is aligned with the grating’s normal direction. As a consequence, we have that *p*(***m***_*i*_|***υ***) = Normal(***m***_*i*_; *U_i_* · ***υ***, *Q_i_*).

Perception of a single grating is known to be affected by contrast. We assume that the noise variance is proportional to the inverse power of the relative contrast *c_i_* = *C_i_*/*C*, i.e. 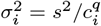, where *q* > 0 is the power exponent and *s*^2^ is the variance corresponding to a relative contrast *c_i_* = 1. This model is consistent with the findings of (24), who derived a similar expression. This expression also predicts that zero contrast (i.e., no grating) corresponds to an infinite noise variance. As a consequence, the covariance matrix of each grating is a function of the contrast: *Q_i_* = *Q_i_*(*c_i_*).

#### Bayesian generative model of plaid perception

We used a Bayesian framework to model the way humans perceive plaid motion (24, 26, 44). The optimal estimate of plaid velocity, ***υ***, from the observed gratings velocities, ***m***_1_ and ***m***_2_, is the one which maximizes the posterior probability of ***υ***, given ***m***_1_ and ***m***_2_:

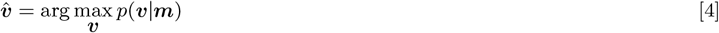

From Bayes’ theorem, the posterior probability is given by: *p*(***υ***|***m***_1_, ***m***_2_) ∝ *L*(***υ***) · *p*(***υ***), where *L*(***υ***) = *p*(***m***_1_|***υ***) · *p*(***m***_2_|***υ***) is the likelihood of ***υ*** given the observations (***m***_1_ and ***m***_2_), whereas *p*(***υ***) is the velocity prior, which reflects prior experience of the subject with observation of moving stimuli. Several studies have reported a perceptual bias toward low-velocity stimuli, which was modeled as a zero-mean, exponential (44) or power-law (26) probability density function. Here, we assume a Gaussian dependence: 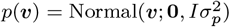.

The above perceptual model implies that the contrasts of the gratings affect plaid velocity estimation through the gratings covariances, *Q_i_*(*c_i_*). In fact, when the two gratings have the same contrast they equally activate the corresponding Fourier (bandpass) motion channels and they equally contribute to the perception of the moving plaid. However, there is some evidence that perceiving the velocity of a single grating is affected by vision of another moving grating with a different contrast (27, 28). Hence in the case of contrast unbalance, i.e. Δ*c* ≠ 0, one grating systematically affects the perception of the other. To incorporate this effect, we tentatively assumed that the perceptual system uses an inaccurate representation of plaid geometry, 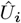, thus generating inaccurate predictions of the grating velocities. We specifically set 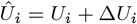, where Δ*U*_1_ = *kU*_2_ Δ*c* and similarly Δ*U*_2_ = *kU*_1_ Δ*c*, in which *k* denotes the amount of cross-talk. A consequence of this inaccurate representation of plaid geometry is that each grating is perceived as slightly rotated toward the other, in a way that is proportional to the relative contrast difference. In conclusion, our Bayesian perceptual model assumes that contrast unbalance has both a systematic and a random effect (on *U_i_* and *Q_i_, i* = 1, 2, respectively).

The optimal estimate of plaid velocity, 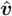 is a random variable (different ***m***_*i*_’s give a different estimate) with a normal distribution, in which both mean and covariance depend on the relative contrast difference, 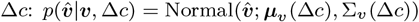. Notice that, because of the prior, the estimate is biased — i.e., the estimator’s expected value is not the true plaid velocity ***υ***. Different from earlier Bayesian formulations (25, 26), the proposed model predicts two key empirical findings about the error in perceived plaid direction: (i) the error decreases with the logarithm of the contrast ratio (27) and (ii) the error is directed towards the higher contrast grating at high plaid speeds, but when the speed decreases the perceived plaid direction is biased towards the low contrast grating (28); see the SI Appendix for details.

#### Perceptual judgement task

The probability of estimating a plaid direction 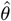 given a specific Δ*c* is given by

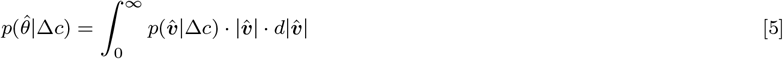

The perceptual judgement task can be modelled as a binary decision between two possible answers, Test (T) or Reference (R). The probability of answering T as a function of the contrast difference Δ*c_T_* in the Test stimulus and Δ*c_R_* in the Reference stimulus, i.e. 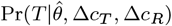, where 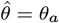 (45 deg in our experiment), can be calculated from Bayes’ theorem:

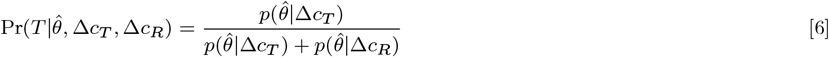

Note that the model predicts that for Δ*c_T_* = Δ*c_R_*, the posterior probability is 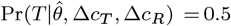. By decreasing Δ*c_T_*, the 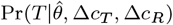 is expected to be greater. Hence for a given value of Δ*c_R_* and 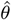, the function 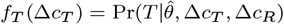 can be interpreted as a psychometric curve, whose magnitude ranges between 0.5 and 1.

Figure 4 summarizes the proposed Bayesian model of plaid perception. Relative contrast modulates the distribution of the estimated plaid velocity. Two relative contrast conditions, one fixed (*R*) and one variable (*T*), are used to build a psychometric curve which denotes the probability of selecting plaid T when asked which of plaid T or R has a movement direction which is closest to the displayed cue.

#### Estimation of model parameters

The psychometric curve of Eq. 6 is a function of the model parameters 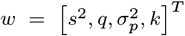, i.e. 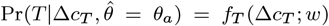. We identified the model parameters *w* from the perceptual judgement data before and after each of the training conditions. The available dataset, 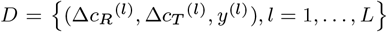, was obtained from repeated forced-choice tests with different values of 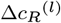 and 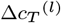, where *y*^(*l*)^ is the T/R answer to the *l*-th test trial (we assume that if T is chosen then *y*^(*l*)^ = 1; *y*^(*l*)^ = 0 otherwise). The answer *y* can be modeled as a random variable with a binomial distribution: Pr(*y*) = *p^y^* · (1 – *p*)^1−*y*^, where 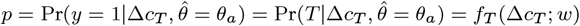.

The optimal estimate of the model parameters given the data were obtained by maximizing the model log-likelihood, assuming that the *L* trials of the perceptual task are independent. The likelihood is given by:

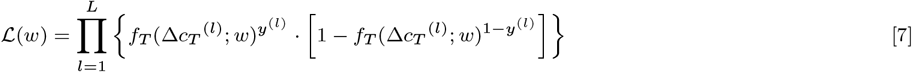

For each subject and for each condition (before and after training), we estimated the model parameters *w* through numeric maximization of 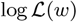.

## ACKNOWLEDGMENTS

This research was supported by a grant from the National Institute of Child Health and Human Development (NICHD R01 HD075740) and a grant from the Italian Ministry of Education, University and Research – Research Projects of National Interest (PRIN 2015).

## References

1. R Held, A Hein, Movement-produced stimulation in the development of visually guided behavior. J. comparative physiological psychology 56, 872 (1963).

2. DJ Ostry, M Darainy, AA Mattar, J Wong, PL Gribble, Somatosensory plasticity and motor learning. J. Neurosci. 30, 5384–5393 (2010).

3. S Vahdat, M Darainy, TE Milner, DJ Ostry, Functionally specific changes in resting-state sensorimotor networks after motor learning. J. Neurosci. 31, 16907–16915 (2011).

4. AA Mattar, M Darainy, DJ Ostry, Motor learning and its sensory effects: time course of perceptual change and its presence with gradual introduction of load. J. neurophysiology 109, 782–791 (2012).

5. EK Cressman, DY Henriques, Sensory recalibration of hand position following visuomotor adaptation. J. neurophysiology 102, 3505–3518 (2009).

6. R Volcic, C Fantoni, C Caudek, JA Assad, F Domini, Visuomotor adaptation changes stereoscopic depth perception and tactile discrimination. J. Neurosci. 33, 17081–17088 (2013).

7. CS Harris, Adaptation to displaced vision: visual, motor, or proprioceptive change? Science 140, 812–813 (1963).

8. PA Beckett, Development of the third component in prism adaptation: effects of active and passive movement. J. Exp. Psychol. Hum. Percept. Perform. 6, 433 (1980).

9. JL Mirdamadi, HJ Block, Somatosensory changes associated with motor skill learning. J. Neurophysiol. 123, 1052–1062 (2020).

10. AV Cuppone, M Semprini, J Konczak, Consolidation of human somatosensory memory during motor learning. Behav. brain research 347, 184–192 (2018).

11. S Schütz-Bosbach, W Prinz, Perceptual resonance: action-induced modulation of perception. Trends Cogn Sci 11, 349–55 (2007).

12. J Zwickel, M Grosjean, W Prinz, Seeing while moving: measuring the online influence of action on perception. Q J Exp Psychol (Hove) 60, 1063–71 (2007).

13. A Wohlschläger, Mental object rotation and the planning of hand movements. Percept. & psychophysics 63, 709–718 (2001).

14. P Veto, M Uhlig, NF Troje, W Einhäuser, Cognition modulates action-to-perception transfer in ambiguous perception. J Vis 18, 5 (2018).

15. A Wohlschläger, Visual motion priming by invisible actions. Vis. Res 40, 925–30 (2000).

16. H Wallach, Über visuell wahrgenommene bewegungsrichtung. Psychol. Forschung 20, 325–380 (1935).

17. CL Fennema, WB Thompson, Velocity determination in scenes containing several moving objects. Comput. graphics image processing 9, 301–315 (1979).

18. EH Adelson, JA Movshon, Phenomenal coherence of moving visual patterns. Nature 300, 523 (1982).

19. GR Stoner, TD Albright, VS Ramachandran, Transparency and coherence in human motion perception. Nature 344, 153–5 (1990).

20. J Kim, HR Wilson, Dependence of plaid motion coherence on component grating directions. Vis. Res. 33, 2479–2489 (1993).

21. JM Hupé, N Rubin, The oblique plaid effect. Vis. Res. 44, 489–500 (2004).

22. GR Stoner, TD Albright, Motion coherency rules are form-cue invariant. Vis. Res. 32, 465–475 (1992).

23. LL Kontsevich, CW Tyler, Bayesian adaptive estimation of psychometric slope and threshold. Vis. Res 39, 2729–37 (1999).

24. F Hürlimann, DC Kiper, M Carandini, Testing the bayesian model of perceived speed. Vis. Res 42, 2253–7 (2002).

25. Y Weiss, EP Simoncelli, EH Adelson, Motion illusions as optimal percepts. Nat Neurosci 5, 598–604 (2002).

26. JH Hedges, AA Stocker, EP Simoncelli, Optimal inference explains the perceptual coherence of visual motion stimuli. J. vision 11, 14–14 (2011).

27. LS Stone, AB Watson, JB Mulligan, Effect of contrast on the perceived direction of a moving plaid. Vis. Res 30, 1049–67 (1990).

28. RA Champion, ST Hammett, PG Thompson, Perceived direction of plaid motion is not predicted by component speeds. Vis. Res 47, 375–83 (2007).

29. LE Brown, ET Wilson, MA Goodale, PL Gribble, Motor force field learning influences visual processing of target motion. J. Neurosci. 27, 9975–9983 (2007).

30. IAM Beets, F Rösler, K Fiehler, Nonvisual motor learning improves visual motion perception: evidence from violating the two-thirds power law. J Neurophysiol 104, 1612–24 (2010).

31. K Sung, WT Wojtach, D Purves, An empirical explanation of aperture effects. Proc. Natl. Acad. Sci. 106, 298–303 (2009).

32. D Purves, BB Monson, J Sundararajan, WT Wojtach, How biological vision succeeds in the physical world. Proc. Natl. Acad. Sci. 111, 4750–4755 (2014).

33. A Haith, CP Jackson, RC Miall, S Vijayakumar, Unifying the sensory and motor components of sensorimotor adaptation in *Advances in Neural Information Processing Systems*. pp. 593–600 (2009).

34. JW Krakauer, P Mazzoni, Human sensorimotor learning: adaptation, skill, and beyond. Curr. opinion neurobiology 21, 636–644 (2011).

35. TD Albright, GR Stoner, Visual motion perception. Proc Natl Acad Sci U S A 92, 2433–40 (1995).

36. C Law, J Gold, Neural correlates of perceptual learning in a sensory-motor, but not a sensory, cortical area. Nat Neurosci 11, 505–513 (2008).

37. KW Latimer, JL Yates, MLR Meister, AC Huk, JW Pillow, Single-trial spike trains in parietal cortex reveal discrete steps during decision-making. Science 349, 184–187 (2015).

38. R Zhang, D Tadin, Disentangling locus of perceptual learning in the visual hierarchy of motion processing. Sci. Reports 9, 1557 (2019).

39. VP Ferrera, HR Wilson, Perceived direction of moving two-dimensional patterns. Vis. Res 30, 273–87 (1990).

40. SJ Cropper, KT Mullen, DR Badcock, Motion coherence across different chromatic axes. Vis. Res 36, 2475–88 (1996).

41. DH Brainard, The psychophysics toolbox. Spat Vis 10, 433–6 (1997).

42. M Kleiner, D Brainard, D Pelli, What is new in psychophysics toolbox. Perception 36 (2007).

43. N Prins, The psi-marginal adaptive method: How to give nuisance parameters the attention they deserve (no more, no less). J Vis 13, 3 (2013).

44. AA Stocker, EP Simoncelli, Noise characteristics and prior expectations in human visual speed perception. Nat Neurosci 9, 578–85 (2006).

