## Supplemental Information for "Self-operated stimuli improve subsequent visual motion processing"

In order to provide additional support for the proposed Bayesian model of plaid perception here we compare the model predictions with two sets of empirical observations about the way perception of plaid direction is affected by absolute and relative contrast (1) and by the grating speeds (2). The comparison will be also used to highlight the role of the individual model parameters in the estimated plaid direction.

#### Model summary

The Bayesian model of plaid perception assumes that perception of a single grating is affected by its contrast. The model also assumes a Gaussian plaid velocity prior, with variance $\sigma_p^2$, and a cross-talk parameter, $k$, which accounts for the conjecture that, in a plaid stimulus, the perception of the direction of each individual grating is biased toward the direction of the other.

#### Effect of gratings contrast on perception of direction

We compared the model predictions with the findings of Stone et al (1). They used a plaid moving upwards ($\theta$ = 90 deg) at 2 deg/s and consisting of two gratings, moving in directions $\theta_1$ = 90 - 60 = 30 deg and $\theta_2$ = 90 + 60 = 150 deg with equal velocities (1 deg/s). In their experiment, they used different values of total contrast, $C$: $\{5\%, 10\%, 20\%, 40\%\}$ and for each $C$ they varied the contrast ratio $a = c_1/c_2$. For each combination of $C$ and $a$, they looked at the perceptual bias between the perceived and actual plaid direction, $\Delta\theta = \hat{\theta} - \theta_1$. They found that the perception of the plaid direction is biased toward the higher contrast grating. As a consequence, the bias increases with both the contrast ratio and decreases with total contrast.

We used the model to simulate the same experiment. The model predicts the probability density function of plaid direction as a function of the relative contrast of the two gratings (Eq. 5, main paper) which is reported here:

$$p(\hat{\theta}|\Delta c) = \int_0^{\infty} p(\hat{\boldsymbol{v}}|\Delta c) \cdot |\hat{\boldsymbol{v}}| \cdot d|\hat{\boldsymbol{v}}| \qquad [1]$$

From this expression it is possible to derive the mean and variance of the estimated plaid direction, $\hat{\theta}$. In our simulations we set $q$=2.5, 3 and 3.5, $\sigma_P^2$ = 1, 4, and 9 deg$^2$/s$^2$ and $k$=0 (no cross-talk) and 0.15. We also assumed that the noise variance of a grating at a maximum contrast is inversely proportional to the total contrast, I.e. $s^2 = h/C$. We set $h = 3 \cdot 10^{-3}$ deg$^2$/s$^2$. These values are similar to the estimates obtained from our data (main paper, Figure 4).

The simulation results are summarized in Figure 1, which has exactly the same format as the figures in (1). In particular, the model correctly captures the dependence of direction bias on contrast ratio. When the contrast ratio is greater than 1 – grating 1 has greater contrast than grating 2 – the perceived direction is biased toward grating 1, and vice versa; see Figure 1 (top). The bias also depends on total contrast (greater contrast, lower bias); see Figure 1 (bottom).

The model also predicts that the contrast dependence of the direction bias is increased by a lower prior variance ($\sigma_P^2$) and by a greater power exponent ($q$). In contrast, the effect decreases when cross-talk ($k$) is present.

#### Effect of speed on perceived plaid direction

The work of Champion et al (2) complements the previous study. Specifically, they assessed how the perceived direction of a plaid which is composed of gratings with different contrasts is affected by the gratings' speed. They reported that the perceived plaid direction is biased toward the direction of the high-contrast grating. However, at low speeds (around 1 deg/s) the effect tends to reverse (bias toward the low-contrast grating). This result is in contrast with (1) who used low speeds but found no such reversal.

We used our model to simulate Experiment 2 in their study. As in their study, we used a high contrast grating ($C_1$=60 %) with $\theta_1$ = 90 - 45° and a low contrast grating ($C_2$ = 30 %) with $\theta_2$ = 90 + 45°. This corresponds to a total contrast $C$ = 90 % (greater than in (1)) and to a relative contrast difference $\Delta c$ = 0.33. As in the previous section, we used Eq. 1 to provide the most likely estimate of plaid direction. As in the previous section, in our simulations we set $q$=2.5, 3 and 3.5, $\sigma_P^2$ = 1, 4, and 9 deg$^2$/s$^2$ and $k$=0 (no cross-talk) and 0.15. As in the previous section, we set $h = 3 \cdot 10^{-3}$ deg$^2$/s$^2$. The simulation results are summarized in Figure 2 (black).

These simulations predict an almost zero directional bias over the whole range of speeds. This is indeed consistent with (1), who found that the directional bias tends to dissappear as the total contrast increases. Nevertheless, if in the model the noise variance of the individual gratings is increased ($h = 6 \cdot 10^{-2}$ deg$^2$/s$^2$) – Figure 2 (red trace), the model predictions closely resemble the findings of (2).

Looking at the roles of the individual model parameters in determining this finding, bias reversal seems to require a combination of higher power exponent ($q$) in the model of grating noise variance, and higher prior variance ($\sigma_P^2$), corresponding to a greater role of the gratings' likelihood in comparison with the velocity prior.

### 1. Conclusions

These results suggest that, in contrast with the conclusions of (2), their results are consistent with a Bayesian model of plaid perception. It should be noted, however, that the two studies use very different contrast values ($C$=90 % vs $C$=5-40 %).

1. LS Stone, AB Watson, JB Mulligan, Effect of contrast on the perceived direction of a moving plaid. *Vis. Res* **30**, 1049–67 (1990).
2. RA Champion, ST Hammett, PG Thompson, Perceived direction of plaid motion is not predicted by component speeds. *Vis. Res* **47**, 375–83 (2007).

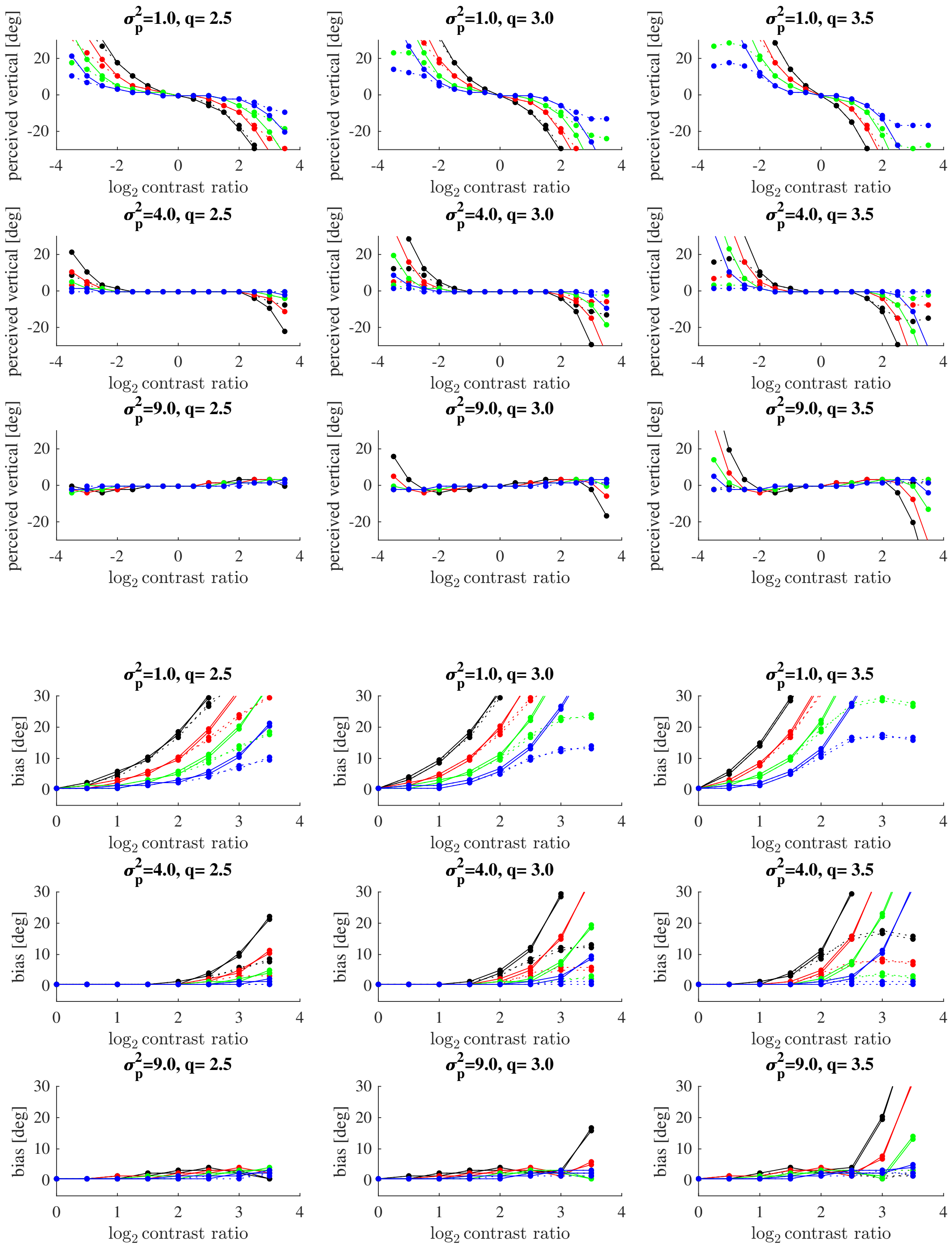


**Fig. 1.** Predicted effect of contrast on perception of plaid direction (1). Perceived vertical (top) and bias (bottom) as functions of total contrast and contrast ratio. The colors refer to different total contrast magnitudes: C=5 % (black), 10 % (red), 20 % (green) and 40 % (blue). The different panels refer to different combinations of power exponent ($q$) and variance of the velocity prior ($\sigma_p^2$). Continuous and dashed lines refer to different amounts of cross-talk ($k$): 0 (continuous) and 0.15 (dashed).

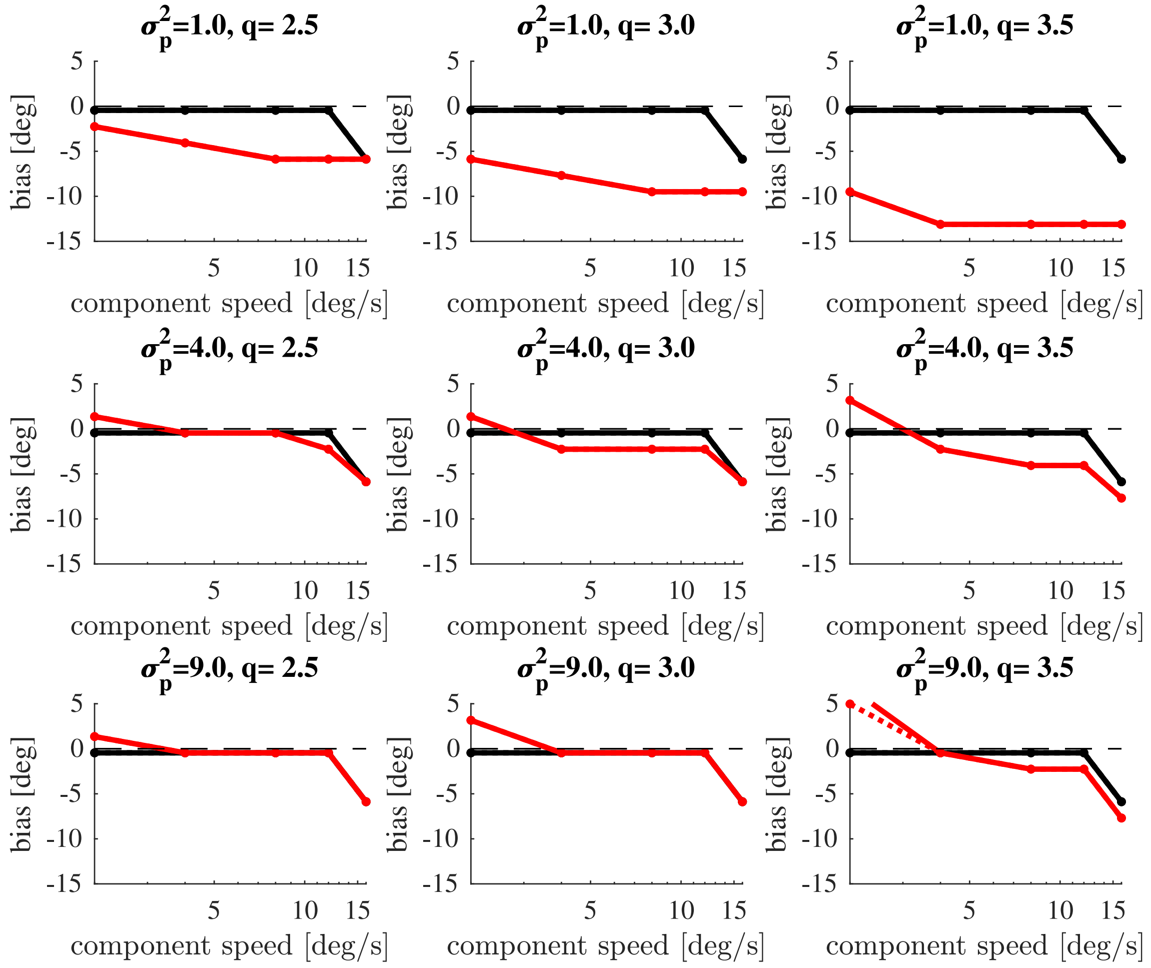


**Fig. 2.** Dependence of the directional bias on the component (grating) speed as in Experiment 2 in (2). The perceived plaid direction is biased toward the high-contrast grating (negative bias) at greater speeds, but the effect tends to reverse (bias toward low-contrast grating) at lower speeds. Black and red lines correspond, respectively, to $h = 3 \cdot 10^{-3}$ deg$^2$/s$^2$ and $h = 6 \cdot 10^{-2}$ deg$^2$/s$^2$. Continuous and dashed lines refer to different amounts of cross-talk ($k$): 0 (continuous) and 0.15 (dashed).
